# Genetic polymorphism of Cytochrome-P450-2C9 (CYP2C9) in Indian populations

**DOI:** 10.1101/113993

**Authors:** Sheikh Nizamuddin, Shivendra Dubey, Sakshi Singh, Saurav Sharma, Anshuman Mishra, K Harish, Harsh Joshi, K. Thangaraj

## Abstract

Cytochrome-P450-2C9 (*CYP2C9*) metabolizes wide range of drugs and highly express in human liver. Various mutations of *CYP2C9* (R144C, I359L etc.), associated with drug-response, are highly diverse. We aimed to investigate the genetic diversity of *CYP2C9* in Indian-subcontinent, using 1278 subjects from 36 populations. High frequency of *CYP2C9*3* (0-0.179) was observed, comparative to other populations, including Europeans. Subjects having *CYP2C9*3/*3* requires lower dose of warfarin, comparative to *CYP2C9*1/*3* or *CYP2C9*1/*1*. Since, Indians are practicing marriage among their caste system, we predicted and observed high frequency (0-0.05) of *CYP2C9*3/*3*. Out of 21 populations, living outside of Indian subcontinent, only Toscani and Southern Han-Chinese have 0.009 and 0.01 *CYP2C9*3/*3*, respectively, lower than Indians,. We found a non-synonymous mutation (*L362V*), observed only in Indian-subcontinent, and have 0-0.056 allelic, 0-0.037 *L/V* and 0-0.037 *V/V* genotype frequency. We observed unfavorable interatomic interactions between hydroxylation sites of warfarin and reactive oxyferryl heme in mutant, comparative to wild-type *CYP2C9*, in molecular dynamic simulations; and predict lower kinetic activity.

## Introduction

Heterogeneous drug response is the major hurdle in the successful treatment of diseases and depends on the genetic variations of drug metabolizing enzyme genes. Cytochrome P450 (CYP) family is an important enzyme of ADME (related to absorption, distribution, metabolism and excretion of drug) genes, of which *CYP2C9* is the major constituent of CYP2C subfamily in human liver. It metabolizes wide range of drugs including anticoagulant (warfarin), non-steroidal anti-inflammatory (celecoxib, diclofenac), anti-diabetic (netaglinide, tolbutamide), anti-hypertensive (irbesartan, losartan) and anti-epileptic (phenytoin)^1^. Several variations in *CYP2C9* have been reported which affects metabolism of the drug. Most notable variations are *CYP2C9*2* (R144C) and *CYP2C9*3* (I359L) which decreases 12% and 5% enzyme activity, respectively^2^. Interestingly, these variations are highly heterogeneous among world population; (1) 8-19% and 3.3-16.3% in Caucasian; (2) 0-0.1% and 1.1-3.6% in Asian; (3) 2.9% and 2.0% in African-American; and (4) 0-4.3% and 0-2.3% in Black/African, respectively^3^. Moreover, other rare and functionally relevant variations were also reported in various populations, which includes; (1) *CYP2C9*6*, 0.6% frequency in African-Americans^4^; (2) *CYP2C9*4*, 0.5% in African-Americans and 6% in Caucasians^2, 5^; and (3) *CYP2C9*13*, 0.19-0.45% in Asian^6^. Recently, Dai et al. (2013) reported several rare variants in the Han Chinese population^7^.

Several studies have been performed on *CYP2C9* in Indian populations. However, most of studies focused only on *CYP2C9*3* and *CYP2C9*2*. Recently, Anil et al (2014) found that *CYP2C9*3* present only in Indo-European population with 0.38–1.85%, while absent in Dravidian, Austro-Asiatic and Tibeto-Burman populations^8^. Indian populations are well known for their endogamy practices and must have very high frequency of homozygous allele^9^, however, Anil et al (2014) did not observe any homozygous *CYP2C9*3/*3* genotype. Many studies have shown that the variations in *CYP2C9* are associated with therapeutic heterogeneity in Indian populations. *CYP2C9*2* and *\*3* has been reported with less hydroxylation (or metabolism) of phenytoin *in vivo* in South-Indian populations^10^, comparative to wild type *CYP2C9*1*. Ramasamy et al. (2007) reported phenytoin toxicity in a patient with normal dose of 300mg/day, who had *CYP2C9*3/*3* genotype^11^. The same symptoms were reported by Thakkar, A. N. et al. (2010) in South-Indian populations^12^. Many studies have also demonstrated that the Indian populations need high dose of warfarin and phenytoin compared to Caucasians^12^. Both of these drugs are metabolized by *CYP2C9*. Some of the drugs, metabolized by *CYP2C9* have narrow therapeutic index *e.g*. warfarin, phenytoin and tolbutamide. This is the reason that small change in the activity of *CYP2C9* may cause major changes in individual’s response.

Considering the high genetic diversity in Indian sub-continent, we explore functionally relevant variations in *CYP2C9*, in the present study. The outcome can be utilized to understand heterogeneous therapeutic response and in development of personalized therapy in Indian sub-continent.

## Material and methods

### Details of samples

In total, 1278 samples were selected for the study. To find the distribution of *CYP2C9* allele/genotype frequency within Indian subcontinent, 36 populations of different linguistic groups and geographical locations were selected (**Table S1**)^9, 13^; and further compared with populations of 1000 genome project. Present work has been approved by the Institutional Ethical Committee of CSIR-Centre for Cellular and Molecular Biology (CSIR-CCMB), Hyderabad, India. Informed written consent has been obtained from every participant, prior to collection of blood samples.

### Re-sequencing of CYP2C9, genotyping and analysis

All the 9 exons, their respective intron-exon boundary, 3’ and 5’ UTR of *CYP2C9* have been resequenced. For designing of primer, DNA sequence of ENST00000260682 from Ensembl (v75) has been used. Out of 3 mRNA of *CYP2C9*, only ENST00000260682 translate to protein. Primer3.0 web-based tool (http://simgene.com/Primer3) was used for designing the primers and further primers specificity were checked with NCBI-primer blast. The details of primer sequences are given in **Table S2**. Polymerase chain reaction (PCR) was performed in 10.0 μl solution, which contains 5.0 μl of 2x EmeraldAmp GT PCR master mix, 10-20 ng of genomic DNA and 0.1 pmole (final concentration) of each primer. Thermal cycling conditions used are as follows: initial denaturation step of 5 min at 94°c, followed by 35 cycles of denaturation step of 30 sec at 94°c, annealing step of 30 sec at their respective melting temperature, extension step of 2 min at 72°c, followed by single step of final extension of 7 min at 72°c. PCR products were cleaned with Exo-SAP-IT (USB, Affymetrix, USA) with recommended protocol of the manufacturer. Cleaned PCR product (1.0 μl) has been subjected to sequencing PCR using BigDye terminator (v3.1) cycle sequencing kit (Applied Biosystem, USA) and analyzed using ABI 3730xl DNA sequencer. AutoAssembler (v1.0) was used for assembling and manual editing of sequence data.

### Distal effect of L362V on kinetics of CYP2C9: molecular dynamics simulation

#### Preparation of 3D (3 dimensional) structure

We performed the molecular dynamics simulation with Gromacs (version 5.0.2)^14^. Starting structure of *CYP2C9* was obtained from PDB (code: 1OG5). We removed warfarin drug from 1OG5 with Chimera (version 1.11)^15^ and utilized in the further structure modifications. In total, 7 amino acids (K206E, I215V, C216Y, S220P, P221A, I223L and I224L) were substituted in wild type *CYP2C9*, to enhance the crystallization^16^. Hence, we modified back 1OG5 to wild type sequence with FoldX (version 2.6) and further utilized to generate mutant proteins. We generated 3D structure of L362V, with FoldX.

#### Docking of warfarin drug

Since, warfarin was located farther from heme center and in catalytically non-reactive stage; we performed docking with AutoDock Vina (version 1.1.2) to obtain putative functional confirmation of drug in the active site of *CYP2C9^17^*. Autodock tools were used for generating partial charges of warfarin and *CYP2C9* using Gasteiger method^18^. Confirmation of warfarin, having lowest binding energy, was selected in further analysis.

#### Simulation

To perform molecular dynamics simulation, Gromacs (version 5.0.2) was used^14^. We assigned the partial charges of warfarin and oxyferryl state of heme group, as described by Seifert, A *et. al* (2006)^19^, while for generating partial charges in wild-type/mutant protein, we used Amber force field “ff99SB”. Before assigning partial charges and structure modifications, we removed H atom from ligand bound protein, with Babel software^20^. Further, structure was loaded in Tleap (AmberTools)^21^ to modify; (1) creating single bond S-Fe, (S of CYM406 and Fe of heme group), (2) creating single bond Fe-O (Fe of heme group and O of water molecule/H_2_O464) and (3) removal of H atom from H_2_O464.

For all ligand bound models (wild-type/mutant), initially we placed the molecule in 1nm cubic box and then solvated it with TIP3P water molecules; and neutralize the system with sodium (NA) molecules. Energy minimization of the system was performed in 10,000 steps with steepest descent minimization algorithm. The potential energy of system has been demonstrated in **Figure S1**. Further, we equilibrated the system at ~310K (normal human body temperature) and at pressure ~1 bar in 20 and 100 pico seconds (ps) respectively in 2 different steps (**Figure S2**). After this, system was simulated for 15 nano seconds (ns), in 4 replicates. To calculate the rolling mean in 100 ps, we utilized the zoo package of R^22^.

### Results and discussions

#### Diversity of CYP2C9*3 in Indian populations

The A→C (rs1057910/ *CYP2C9*3)* is a non-synonymous mutation, which replace Isoleucine with Leucine (ATT>CTT; Ile359Leu; low enzyme activity). Considering the higher level of evidence between *CYP2C9*3* and drug response, CPIC (Clinical Pharmacogenomics Implementation Consortium) has been categorized *CYP2C9*3* under level-1A^23^. *CYP2C9*3* has been reported with hypersensitive reaction against phenytoin in epilepsy patients^24^, decreased metabolism of celecoxib^25^. It is also reported with high incident of response rate against sulfonamides, urea derivatives^26^.

To explore the “C” allele frequency in Indian populations, initially we confirmed Hardy-Weinberg equilibrium (HWE). It was observed that 10 populations were not in HWE (*p*-value < 0.05), which include 1 Indo-European population, Haryana Pandit (p-value = 1.3×10^−4^) and 9 Dravidians populations; Mudaliar and Madar from Tamil Nadu (p-value = 1.92×10^−6^ and 4.75×10^−7^ respectively), Gawali from Karnataka (p-value = 4.12×10^−4^), Kurumba from Kerala (p-value = 6.94×10^−6^) and Telagas, Thoti, Chenchu, Patkar and Vaddera from Andhra Pradesh (p-value = 1.6×10^−3^, 2.23×10^−8^, 0, 5.21×10^−6^ and 6.99×10^−3^ respectively) (Table 1).

**Table 1.** Distribution of CYP2C9*3 (I359L) in Indian populations

| Populations | State | Lat. | Long. | Linguistic | Sample size | Genotype/Alele percentage (number of individuals): rs1057910 |  |  |  |  |  | HWE p-value |
| --- | --- | --- | --- | --- | --- | --- | --- | --- | --- | --- | --- | --- |
|  |  |  |  |  |  | Missing data | AA | AC | CC | A | C |  |
| Mahli | Jharkhand | 23.46 | 85 | Austro-Asiatic | 38 | 4(10.526) | 27(79.412) | 6(17.647) | 1(2.941) | 60(88.235) | 8(11.765) | 0.3712145 |
| Gond | Chattisgarh | 19.87 | 81.6 | Austro-Asiatics | 37 | 8(21.622) | 28(96.552) | 0(0) | 1(3.448) | 56(96.552) | 2(3.448) | 0.018 |
| Kharia | Chattisgarh | 23.33 | 85.44 | Austro-Asiatics | 86 | 14(16.279) | 62(86.111) | 10(13.889) | 0(0) | 134(93.056) | 10(6.944) | 1 |
| Gond | Madhya-Pradesh | 26.12 | 77.4 | Austro-Asiatics | 38 | 7(18.421) | 25(80.645) | 6(19.355) | 0(0) | 56(90.323) | 6(9.677) | 1 |
| Ho | Jharkhand | 23.35 | 85.33 | Austro-Asiatics | 67 | 2(2.985) | 47(72.308) | 16(24.615) | 2(3.077) | 110(84.615) | 20(15.385) | 0.633 |
| Kolhas | Andhra-Pradesh | 14.46 | 79.98 | Dravidian | 14 | 2(14.286) | 10(83.333) | 2(16.667) | 0(0) | 22(91.667) | 2(8.333) | 1 |
| Adi-Dravidar | Tamilnadu | 11.35 | 77.73 | Dravidian | 15 | 1(6.667) | 9(64.286) | 5(35.714) | 0(0) | 23(82.143) | 5(17.857) | 1 |
| Telagas | Andhra-Pradesh | 18.17 | 83.53 | Dravidians | 12 | 0(0) | 9(75) | 0(0) | 3(25) | 18(75) | 6(25) | 0.001634521 |
| Thoti | Andhra-Pradesh | 16.51 | 80.64 | Dravidians | 29 | 0(0) | 20(68.966) | 0(0) | 9(31.034) | 40(68.966) | 18(31.034) | 2.23E-08 |
| Naidu | Andhra-Pradesh | 13.22 | 79.6 | Dravidians | 21 | 11(52.381) | 9(90) | 1(10) | 0(0) | 19(95) | 1(5) | 1 |
| Reddy | Andhra-Pradesh | 17.37 | 78.48 | Dravidians | 24 | 1(4.167) | 17(73.913) | 6(26.087) | 0(0) | 40(86.957) | 6(13.043) | 1 |
| Mudaliar | Tamilnadu | 12.92 | 79.13 | Dravidians | 48 | 3(6.25) | 41(91.111) | 0(0) | 4(8.889) | 82(91.111) | 8(8.889) | 1.92E-06 |
| Gammavokklu | Karnataka | 12.93 | 74.83 | Dravidians | 19 | 4(21.053) | 13(86.667) | 2(13.333) | 0(0) | 28(93.333) | 2(6.667) | 1 |
| Vysya | Andhra-Pradesh | 14.68 | 77.65 | Dravidians | 60 | 10(16.667) | 40(80) | 10(20) | 0(0) | 90(90) | 10(10) | 1 |
| Gawli | Karnataka | 13.33 | 74.77 | Dravidians | 89 | 10(11.236) | 63(79.747) | 10(12.658) | 6(7.595) | 136(86.076) | 22(13.924) | 0.000412713 |
| Medari | Andhra-Pradesh | 16.56 | 80.61 | Dravidians | 4 | 0(0) | 3(75) | 1(25) | 0(0) | 7(87.5) | 1(12.5) | 1 |
| Madar | Karnataka | 15.33 | 75.05 | Dravidians | 70 | 9(12.857) | 55(90.164) | 1(1.639) | 5(8.197) | 111(90.984) | 11(9.016) | 4.75E-07 |
| Patkar | Andhra-Pradesh | 15.8 | 78.1 | Dravidians | 20 | 1(5) | 12(63.158) | 0(0) | 7(36.842) | 24(63.158) | 14(36.842) | 5.21E-06 |
| Raj-Gond | Madhya-Pradesh | 23.87 | 78.7 | Dravidians | 28 | 19(67.857) | 9(100) | 0(0) | 0(0) | 18(100) | 0(0) | 1 |
| Adhiyan | Tamilnadu | 13.72 | 79.41 | Dravidians | 44 | 4(9.091) | 37(92.5) | 3(7.5) | 0(0) | 77(96.25) | 3(3.75) | 1 |
| Kurumba | Tamilnadu | 12.94 | 79.09 | Dravidians | 15 | 2(13.333) | 11(84.615) | 2(15.385) | 0(0) | 24(92.308) | 2(7.692) | 1 |
| Chenchu | Andhra-Pradesh | 17.37 | 78.47 | Dravidians | 27 | 2(7.407) | 17(68) | 0(0) | 8(32) | 34(68) | 16(32) | 0 |
| Kurumba | Madhya-Pradesh | 22.71 | 75.83 | Dravidians | 26 | 6(23.077) | 14(70) | 0(0) | 6(30) | 28(70) | 12(30) | 6.94E-06 |
| Vaddera | Andhra-Pradesh | 18.72 | 79.48 | Dravidians | 8 | 0(0) | 5(62.5) | 0(0) | 3(37.5) | 10(62.5) | 6(37.5) | 0.006993007 |
| Brahmin-Tiwari | Uttar-Pradesh | 25.73 | 82.68 | Indo-European | 44 | 13(29.545) | 28(90.323) | 3(9.677) | 0(0) | 59(95.161) | 3(4.839) | 1 |
| Kashmiri pandit | JammuKashmir | 34.37 | 75.83 | Indo-European | 21 | 0(0) | 17(80.952) | 3(14.286) | 1(4.762) | 37(88.095) | 5(11.905) | 0.235 |
| Bhil | Gujarat | 23.03 | 72.67 | Indo-European | 4 | 0(0) | 4(100) | 0(0) | 0(0) | 8(100) | 0(0) | 1 |
| Gamit | Gujrat | 21.17 | 72.83 | Indo-European | 45 | 7(15.556) | 35(92.105) | 3(7.895) | 0(0) | 73(96.053) | 3(3.947) | 1 |
| Tharu | Uttarakhand | 29.38 | 79.5 | Indo-European | 30 | 3(10) | 23(85.185) | 3(11.111) | 1(3.704) | 49(90.741) | 5(9.259) | 0.183 |
| Warli | Maharashtra | 19.17 | 72.95 | Indo-European | 70 | 7(10) | 48(76.19) | 15(23.81) | 0(0) | 111(88.095) | 15(11.905) | 0.5832885 |
| Baiswar | Uttar-Pradesh | 25.15 | 82.6 | Indo-Europeans | 40 | 6(15) | 23(67.647) | 11(32.353) | 0(0) | 57(83.824) | 11(16.176) | 0.5620674 |
| Pandit | Haryana | 29.96 | 76.87 | Indo-Europeans | 40 | 12(30) | 23(82.143) | 1(3.571) | 4(14.286) | 47(83.929) | 9(16.071) | 0.000129726 |
| Bhilala | Madhya-Pradesh | 22.6 | 75.3 | Indo-Europeans | 49 | 9(18.367) | 33(82.5) | 5(12.5) | 2(5) | 71(88.75) | 9(11.25) | 0.05670192 |
| Chakhesang_Naga | Nagaland | 26.12 | 94.48 | Tibeto-Burmans | 33 | 19(57.576) | 14(100) | 0(0) | 0(0) | 28(100) | 0(0) | 1 |
| Naga-sema | Nagaland | 25.7 | 93.81 | Tibeto-Burmans | 40 | 21(52.5) | 16(84.211) | 3(15.789) | 0(0) | 35(92.105) | 3(7.895) | 1 |
| Mizo | Mizoram | 23.2 | 92.83 | Tibeto-Burmans | 23 | 7(30.435) | 13(81.25) | 3(18.75) | 0(0) | 29(90.625) | 3(9.375) | 1 |

After excluding these 10 populations, we estimated 11.67% (183 out of 1568) “C” allele in Indian populations, similar (p-value = 0.617) to South-Asian populations of 1000 genome project. Further, we categorized these samples on the basis of their linguists and observed that Dravidians have higher percentage of “C” allele (15.32%; 72 out of 470) while Tibeto-Burman have lowest (6.12%; 6 out of 92). Moreover, in Austro-Asiatics and Indo-European, we observed 9.96% (46 out of 416) and 10.96% (59 out of 538), respectively (Table 1). Interestingly, Tibeto-Burman are insignificantly different (p-value = 0.21) from East-Asians. Adi-Dravidians (schedule tribe) of TamilNadu, Ho (schedule tribe) of Jharkhand and Baiswar (caste) of Uttar-Pradesh have 17.857%, 15.385% and 16.176% of *CYP2C9*3*, respectively, which is higher in their respective linguistic group; while Bhil of Gujarat, Raj-Gond of Madhya-Pradesh and Gond of Chhattisgarh have 0%, 0%, 2%, respectively (Table 1). In Indian sub-continent, high local heterogeneity was observed and any correlation with geographical location does not exist (Figure 1A and Table 1). It is evident in isofrequency map that Indian populations have high frequency of *CYP2C9*3*, comparative to other world populations (Figure 1A). We observed, decreasing gradient of “C” allele frequency from Indian subcontinent to Europeans (Figure 1A).

**Figure 1.**
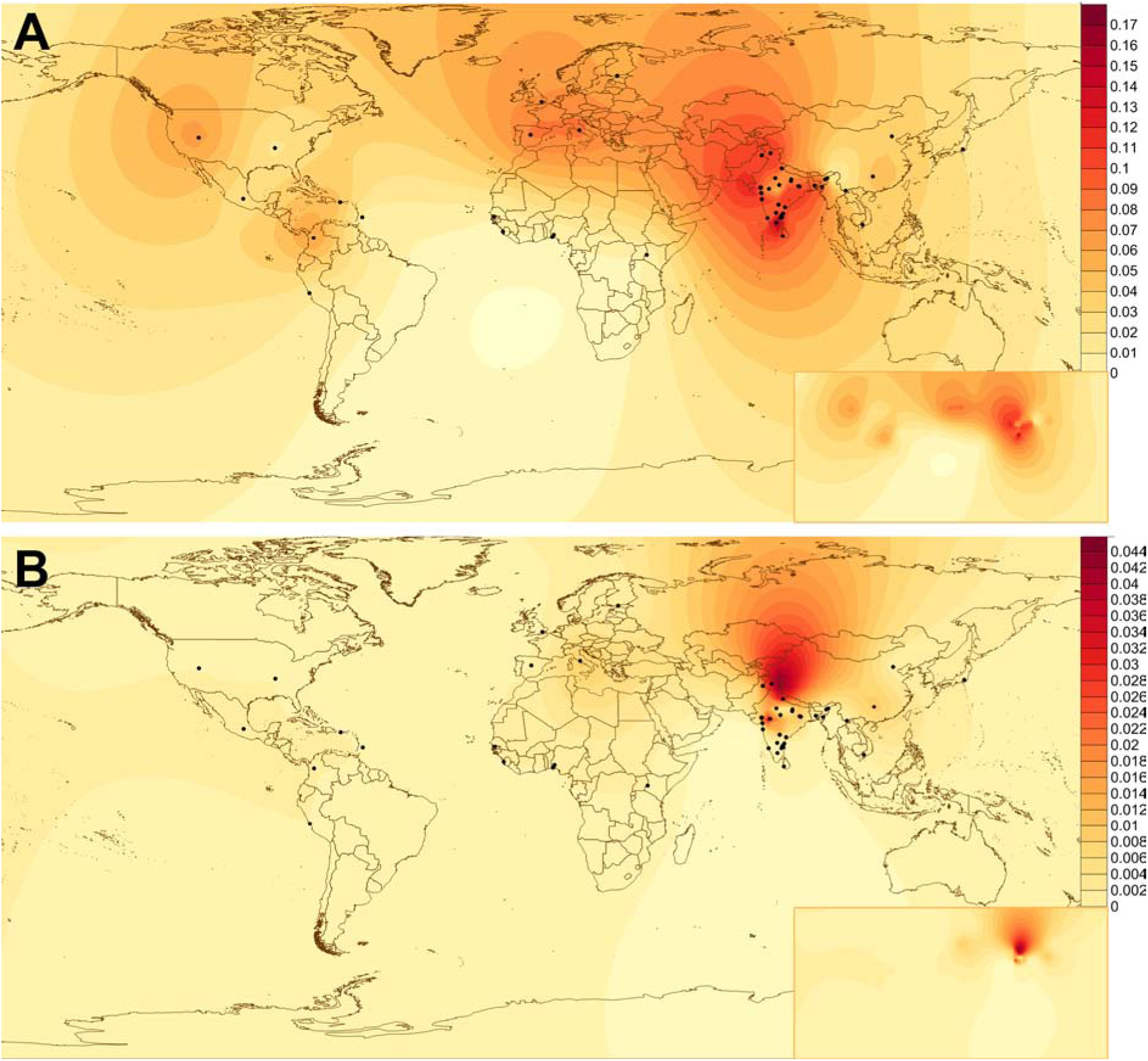
Isofrequency map of (A) *CYP2C9*3* and (B) *CYP2C9*3/*3* to demonstrate the geospatial distribution. We excluded those populations who were not in HWE. In map, dots represent the sampling location.

On the basis of founder events and endogamy marriage practices, we have already predicted high frequency of homozygous alleles in Indian populations^9^. Since, patients with *CYP2C9*3/*3* requires lower dose (0.5-2 mg) of warfarin in comparison to those who have *CYP2C9*1/*3* (3-4 mg), it would be interesting to explore in Indian populations. As expected, we observed higher percentage (<5%) of *CYP2C9*3/*3* in Indians, comparative to other world populations, who have 0-1% (Figure 1B **and** Table 1). Out of 21 populations who are living out of Indian subcontinent, only TSI (Italian populations) and CHS (South Chinese populations) have homozygous genotype (0.9 and 1%), while out of 5 populations who are living in Indian sub-continent, 3 (PJL, ITU and GIH) have 1% *CYP2C9*3/*3*. In present Indian populations samples, we observed 0-5% *CYP2C9*3/*3*, of which Bhilala of Madhya-Pradesh and Ho of Jharkhand have 5% and 3%, respectively; higher in Indo-Europeans and Austro-Asiatic linguistic group (Table 1). We observed 0% *CYP2C9*3/*3* in Tibeto-Burman. Since, *CYP2C9*3/*3* was not in HWE in Dravidian populations, we did not concluded the frequency.

#### Novel non-synonymous variant L362V

We observed a non-synonymous mutation L362V in Indian population at higher frequency. Since, this mutation was observed in higher frequency and present only in Indian populations; we further explored it in 1000 genome project samples. We found that this variant (rs578144976; L362V) was reported at very low frequency (0.4%) in South-Asians, while absent in other world populations. Only ITU (Telugu population) and BEB (Bengali population) are reported to have 0.6% and 1.5% (http://browser.1000genomes.org/Homo_sapiens/Variation/Population?db=core;g=ENSG00000138109;r=10:96698415-96749147;v=ss1338631398;vdb=variation;vf=75731685). In Indian sub-continent, it was observed in all four linguistic groups (13 populations). After excluding the populations in which L362V is not in HWE, we observed 0-5.6% allele, 0-3.7% heterozygous and 0-3.7% homozygous mutant genotype frequency (Table 2, Figure 2A **and** 2B). It can be hypothesize that L362V is originated recently in Indian sub-continent and due to long-term isolation of Indian populations with rest of the world, it was observed only in this region. This might be the reason also, why L362V, deviated from Hardy-Weinberg equilibrium in many populations.

**Figure 2.**
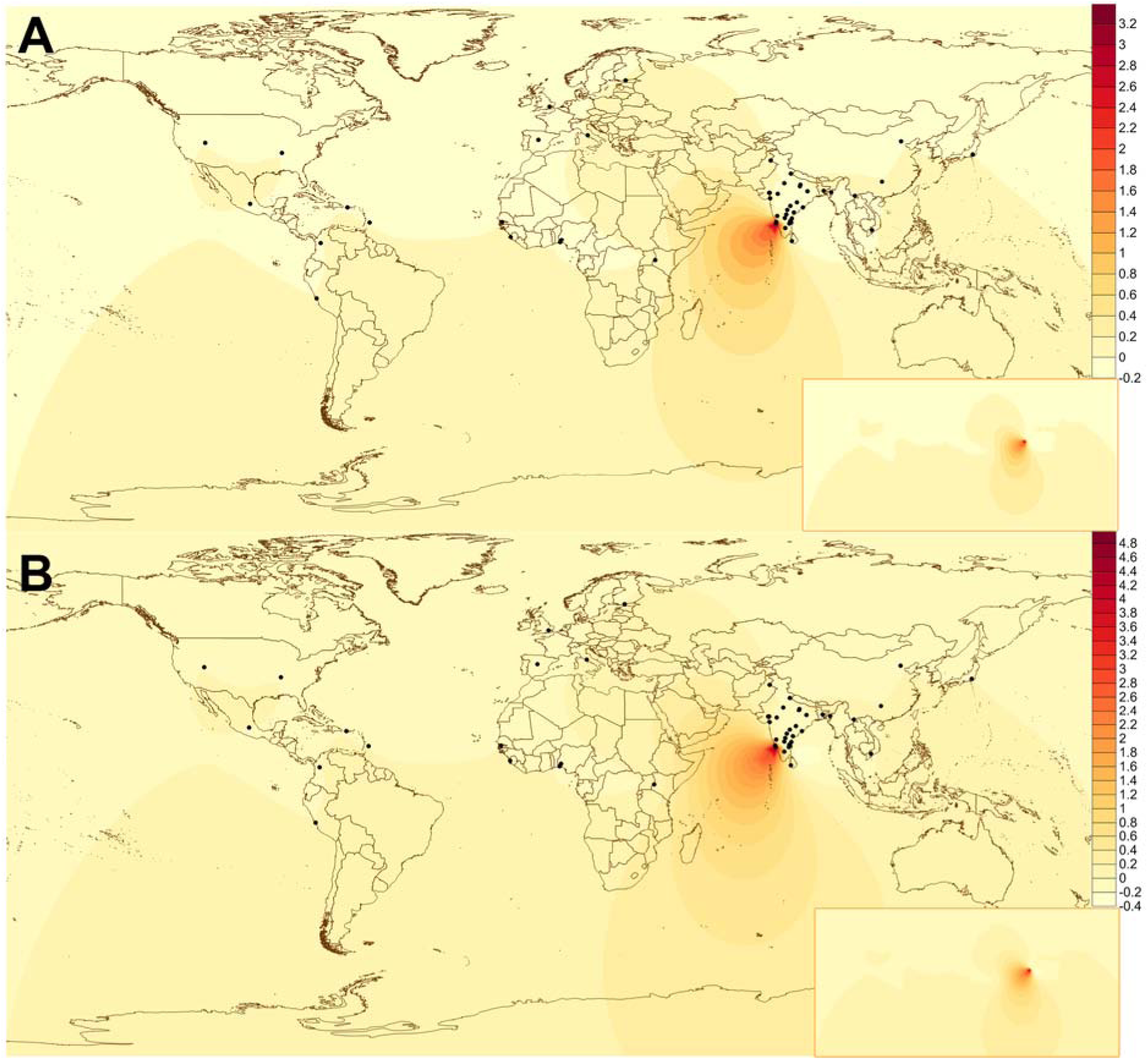
Isofrequency map of novel mutant L362V. Figure (A) and (B) represents the allelic and genotype geospatial distribution. Similar to figure 1, we excluded those populations who were not in HWE.

**Table 2.** Distribution of novel non-synonymous mutation L362V, observed in exon-7 of *CYP2C9*

| Populations | State | Lat. | Long. | Linguistic | Sample size | number of individuals (Genotype/Alelle percentage): Novel; Leu>Val |  |  |  |  |  | HWE P-value |
| --- | --- | --- | --- | --- | --- | --- | --- | --- | --- | --- | --- | --- |
|  |  |  |  |  |  | Missing data | CC | CG | GG | C | G |  |
| Mahli | Jharkhand | 23.46 | 85 | Austro-Asiatic | 38 | 4(21.053) | 15(100) | 0(0) | 0(0) | 30(100) | 0(0) | 1 |
| Ho | Jharkhand | 23.35 | 85.33 | Austro-Asiatics | 67 | 3(6.25) | 41(91.111) | 0(0) | 4(8.889) | 82(91.111) | 8(8.889) | 1.92E-06 |
| Gond | Madhya-Pradesh | 26.12 | 77.4 | Austro-Asiatics | 38 | 11(52.381) | 10(100) | 0(0) | 0(0) | 20(100) | 0(0) | 1 |
| Gond | Chattisgarh | 19.87 | 81.6 | Austro-Asiatics | 37 | 7(18.421) | 31(100) | 0(0) | 0(0) | 62(100) | 0(0) | 1 |
| Kharia | Chattisgarh | 85.44 | 23.33 | Austro-Asiatics | 86 | 13(29.545) | 31(100) | 0(0) | 0(0) | 62(100) | 0(0) | 1 |
| Adi-Dravidar | Tamilnadu | 11.35 | 77.73 | Dravidian | 15 | 7(15.556) | 38(100) | 0(0) | 0(0) | 76(100) | 0(0) | 1 |
| Kolhas | Andhra-Pradesh | 14.46 | 79.98 | Dravidian | 14 | 2(14.286) | 12(100) | 0(0) | 0(0) | 24(100) | 0(0) | 1 |
| Reddy | Andhra-Pradesh | 17.37 | 78.48 | Dravidians | 24 | 6(23.077) | 14(70) | 0(0) | 6(30) | 28(70) | 12(30) | 6.94E-06 |
| Thoti | Andhra-Pradesh | 16.51 | 80.64 | Dravidians | 29 | 12(30) | 23(82.143) | 1(3.571) | 4(14.286) | 47(83.929) | 9(16.071) | 0.000129726 |
| Raj-Gond | Madhya-Pradesh | 23.87 | 78.7 | Dravidians | 28 | 2(2.985) | 63(96.923) | 0(0) | 2(3.077) | 126(96.923) | 4(3.077) | 0.000183117 |
| Chenchu | Andhra-Pradesh | 17.37 | 78.47 | Dravidians | 27 | 9(18.367) | 38(95) | 0(0) | 2(5) | 76(95) | 4(5) | 0.000493178 |
| Kurumba | Madhya-Pradesh | 22.71 | 75.83 | Dravidians | 26 | 9(45) | 9(81.818) | 0(0) | 2(18.182) | 18(81.818) | 4(18.182) | 0.007518797 |
| Kurumba | Tamilnadu | 12.94 | 79.09 | Dravidians | 15 | 0(0) | 20(95.238) | 0(0) | 1(4.762) | 40(95.238) | 2(4.762) | 0.02439024 |
| Gammavokklu | Karnataka | 12.93 | 74.83 | Dravidians | 19 | 3(10) | 25(92.593) | 1(3.704) | 1(3.704) | 51(94.444) | 3(5.556) | 0.05660377 |
| Adhiyan | Tamilnadu | 13.72 | 79.41 | Dravidians | 44 | 2(7.407) | 25(100) | 0(0) | 0(0) | 50(100) | 0(0) | 1 |
| Gawli | Karnataka | 13.33 | 74.77 | Dravidians | 89 | 4(9.091) | 40(100) | 0(0) | 0(0) | 80(100) | 0(0) | 1 |
| Madar | Karnataka | 15.33 | 75.05 | Dravidians | 70 | 0(0) | 12(100) | 0(0) | 0(0) | 24(100) | 0(0) | 1 |
| Medari | Andhra-Pradesh | 16.56 | 80.61 | Dravidians | 4 | 19(67.857) | 9(100) | 0(0) | 0(0) | 18(100) | 0(0) | 1 |
| Mudaliar | Tamilnadu | 12.92 | 79.13 | Dravidians | 48 | 1(5) | 19(100) | 0(0) | 0(0) | 38(100) | 0(0) | 1 |
| Naidu | Andhra-Pradesh | 13.22 | 79.6 | Dravidians | 21 | 10(16.667) | 50(100) | 0(0) | 0(0) | 100(100) | 0(0) | 1 |
| Patkar | Andhra-Pradesh | 15.8 | 78.1 | Dravidians | 20 | 19(57.576) | 14(100) | 0(0) | 0(0) | 28(100) | 0(0) | 1 |
| Telagas | Andhra-Pradesh | 18.17 | 83.53 | Dravidians | 12 | 7(10) | 63(100) | 0(0) | 0(0) | 126(100) | 0(0) | 1 |
| Vaddera | Andhra-Pradesh | 18.72 | 79.48 | Dravidians | 8 | 7(30.435) | 16(100) | 0(0) | 0(0) | 32(100) | 0(0) | 1 |
| Vysya | Andhra-Pradesh | 14.68 | 77.65 | Dravidians | 60 | 1(4.167) | 23(100) | 0(0) | 0(0) | 46(100) | 0(0) | 1 |
| Kashmiri pandit | Jammu Kashmir | 34.37 | 75.83 | Indo-European | 21 | 10(11.236) | 73(92.405) | 0(0) | 6(7.595) | 146(92.405) | 12(7.595) | 8.44E-10 |
| Warli | Maharastra | 19.17 | 72.95 | Indo-European | 70 | 9(12.857) | 56(91.803) | 0(0) | 5(8.197) | 112(91.803) | 10(8.197) | 4.32E-08 |
| Bhil | Gujarat | 23.03 | 72.67 | Indo-European | 4 | 14(16.279) | 72(100) | 0(0) | 0(0) | 144(100) | 0(0) | 1 |
| Brahmin-Tiwari | Uttar-Pradesh | 25.73 | 82.68 | Indo-European | 44 | 0(0) | 4(100) | 0(0) | 0(0) | 8(100) | 0(0) | 1 |
| Gamit | Gujrat | 21.17 | 72.83 | Indo-European | 45 | 1(6.667) | 14(100) | 0(0) | 0(0) | 28(100) | 0(0) | 1 |
| Tharu | Uttarakhand | 29.38 | 79.5 | Indo-European | 30 | 0(0) | 29(100) | 0(0) | 0(0) | 58(100) | 0(0) | 1 |
| Pandit | Haryana | 29.96 | 76.87 | Indo-Europeans | 40 | 1(2.174) | 35(77.778) | 0(0) | 10(22.222) | 70(77.778) | 20(22.222) | 6.26E-11 |
| Baiswar | Uttar-Pradesh | 25.15 | 82.6 | Indo-Europeans | 40 | 2(13.333) | 13(100) | 0(0) | 0(0) | 26(100) | 0(0) | 1 |
| Bhilala | Madhya-Pradesh | 22.6 | 75.3 | Indo-Europeans | 49 | 0(0) | 4(100) | 0(0) | 0(0) | 8(100) | 0(0) | 1 |
| Naga-sema | Nagaland | 25.7 | 93.81 | Tibeto-Burmans | 40 | 4(10.526) | 33(97.059) | 0(0) | 1(2.941) | 66(97.059) | 2(2.941) | 0.01492537 |
| Chakhesang_Naga | Nagaland | 26.12 | 94.48 | Tibeto-Burmans | 33 | 8(21.622) | 28(96.552) | 0(0) | 1(3.448) | 56(96.552) | 2(3.448) | 0.01754386 |
| Mizo | Mizoram | 23.2 | 92.83 | Tibeto-Burmans | 23 | 0(0) | 8(100) | 0(0) | 0(0) | 16(100) | 0(0) | 1 |

#### Effect of L362V: In-Silico prediction

Lertkiatmongkol, P. *et. al* (2013)^27^ predicted distal effect of amino acid substitution in *CYP2C9* in molecular dynamics simulation. Authors observed that distance between hydroxylation sites of warfarin and reactive oxyferryl heme in mutant *(CYP2C9*3*: FeO-C7=4.85 Å; *CYP2C9*13*: FeO-C7=3.56 Å; and *CYP2C9*2*: FeO-C7=4.46 Å) is higher, comparative to wild type protein (FeO-C7=3.30 Å). This unfavorable interatomic interaction is the reason of lower kinetic activity in mutant proteins. We utilized same molecular dynamics simulation to explore the distal effect of L362V on the kinetic activity of *CYP2C9*, comparative to wild type protein.

We performed 4 different simulations for wild type and mutant (L362V) *CYP2C9*, after re-docking warfarin in active site (Figure 3) and equilibrating system (**Figure S2**). The system contains water molecular and wild-type/mutant *CYP2C9* in 1×1×1 nm^3^ cubic box, neutralized by sodium ion (Na^+^). We observed that in all 4 simulations, wild-type and mutant *CYP2C9* protein were in stable state [radius of gyration (R_g_); for wild type, 1^st^: 2.249±0.0062, 2^nd^: 2.246±0.0058, 3^rd^: 2.243±0.0058 and 4^th^: 2.247±0.0064; for mutant, 1^st^: 2.266±0.0104, 2^nd^: 2.263± 0.0062, 3^rd^: 2.59±0.0062 and 4^th^: 2.253±0.0058] (**Figure S3**). Hydroxylation at C6, C7 and C4 of warfarin needs the hydrogen bond between H6, H7 and H4 of warfarin and FeO (oxygen) of oxyferryl heme. Therefore, we consider only those confirmations in active state, which are having rolling mean distance of H^…^O < 3Å (distance of C-H^…^O). We observed unfavorable interatomic interactions for hydroxylation between (1) H6 and FeO and (2) H7 and FeO in simulation of mutant, comparative to wild-type *CYP2C9* (Figure 4, **6** and **S4**). The simulations predict the less kinetic activity of the mutant (L362V) *CYP2C9*, comparative to wild-type.

**Figure 3.**
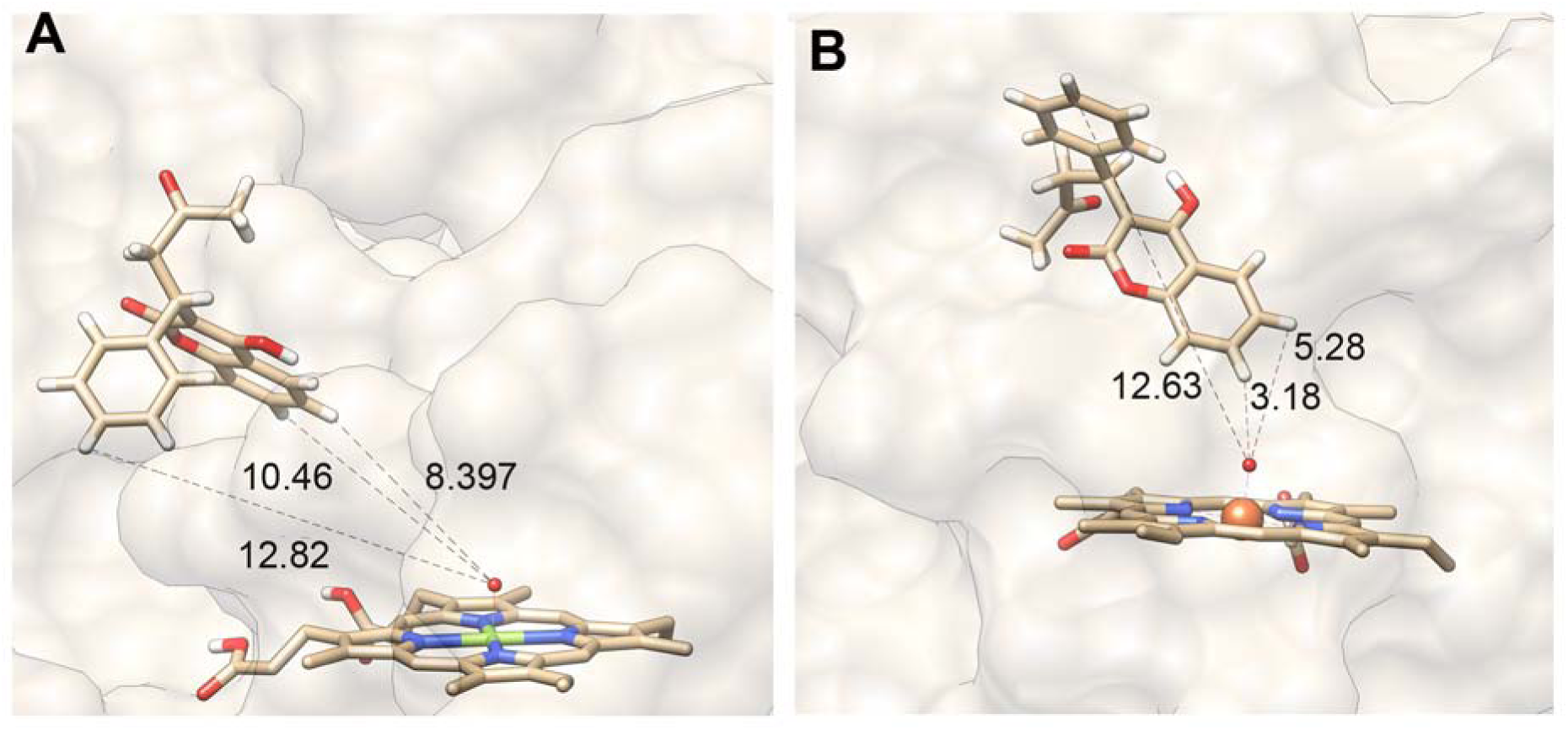
Confirmation of warfarin in (A) original 3D structure (1OG5), dowloaded from PDB and (B) after re-docking.

**Figure 4.**
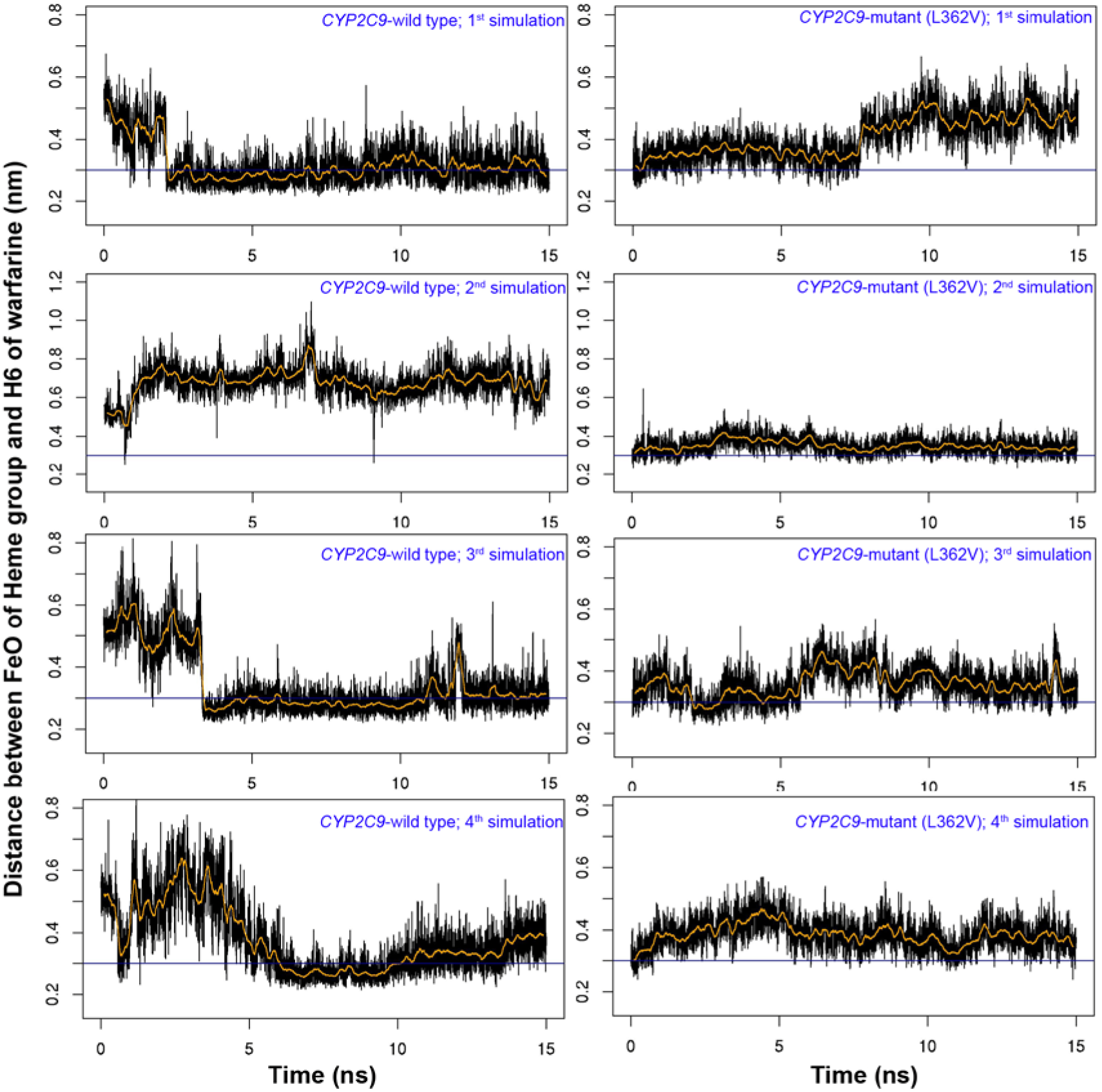
Distance between FeO of oxyferryl heme group and hydrogen atom (H6) bound to C6 of warfarin. The middle orange line represents rolling mean of distance with window size of 100 ps while blue line represents the cut-off value (< 3 Å) for favorable hydrogen bond. Hydrogen atom H6 attains favorable interactions for hydroxylation (H-bond < 3 Å) only in wild-type *CYP2C9* (during 1^st^, 2^nd^ and 3^rd^ molecular dynamics simulation)

**Figure 5.**
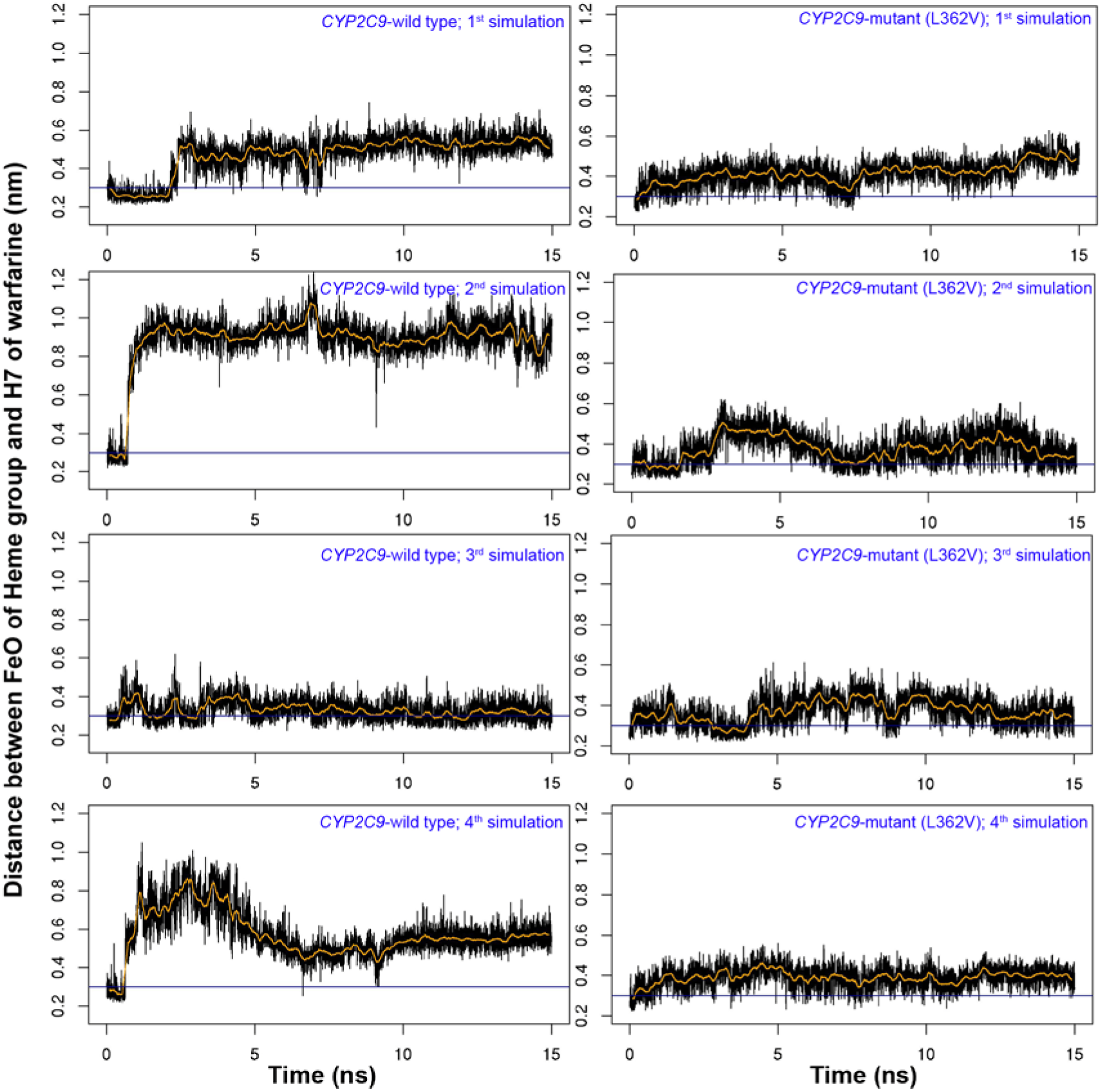
Distance between FeO of oxyferryl heme group and H7 of warfarin. Orange and blue lines represents similar values, given in figure 4. Hydrogen atom H7 attains favorable interactions for hydroxylation (H-bond < 3 Å) only in wild-type *CYP2C9* (during 2^nd^ molecular dynamics simulation).

#### Other rare variants

Besides these, a few rare variants have also been observed in this study. The non-synonymous C>T mutation (rs28371685), which replace arginine with tryptophan (R335W) and determine *CYP2C9*11* haplogroup was found only in 3 individuals (0.2347%, 1 each in Chenchu, Telagas of Andhra Pradesh and Mudliar of Tamil Nadu). Besides this, 2 novel non-synonymous mutations F482L and T283S were found in 2 individual (0.0782%), each in Mizo (Mizoram) and Warli (Maharashtra), respectively.

## Acknowledgements

This work was supported by CSIR Network project—EpiHeD (BSC0118), Government of India. Sheikh Nizamuddin was supported by ICMR JRF-SRF research fellowship. We acknowledge Dr. Gyaneshwar chaubey, for kindly preparing the isofrequency map with Surfer software (version 10). We also acknowledge Dr. Ravi Kumar Verma, who helped us in molecular dynamics simulation.

